# *In-silico* predictive identification of K-Ras^G12V^ inhibitors in natural compounds

**DOI:** 10.1101/553149

**Authors:** Masoud Aliyar, Hassan Aryapour, Majid Mahdavi

## Abstract

As RAS protein is highly significant in signaling pathways, involving cell growth, differentiation and apoptosis; the Ras GTPase proteins play a significant as a master switch in the appearance of many diseases, including 20-30% of all cancers. So, the K-RasG12V mutant was selected as a drug target in present study. This mutant is involved in gastric cancer, lung and pancreatic carcinoma, and colon cancers. So, we employed the structure-based drug design methods and molecular dynamics simulations to undergo virtual screening on natural products small molecules and predicted some new potent therapeutic inhibitors. Finally, ZINC15671852, ZINC85592862, ZINC85567582 and ZINC03616630 final Hits were identified as potent inhibitors from among more than 79,000 bioactive compounds from natural resource. Molecular Mechanics Poisson-Boltzmann Surface Area (MM-P/GBSA) calculation results have also demonstrated that these molecules obtained higher binding free energy than co-crystalized reference ligand.

## Introduction

Ras oncogenes belong to the Ras superfamily of small GTPase proteins, which act as master switch (Fig1) and are involved in intracellular signaling pathways such as cell growth, differentiation and apoptosis [1]. These onco-proteins act as molecular switches, cycling between active (GTP-bound) and inactive (GDP-bound) states. These status changes occur through some conformational changes in two important regions of Ras protein’s surface, called switch-1 (residues 32-38) and switch-2 (Residues 60-75) that are catalyzed via interactions with GEF and GAP factors and the intrinsic GTPase activity [2,3]. Some point mutations, appearing in codons 12, 13 and 61 are significantly responsible for keeping the Ras protein in an active state and have continuous interaction with downstream effectors; such as Raf kinases [4]. These mutated oncogenes are identified and observed in 20-30% of all human cancers including 63-90%, 30-50% and 43% of pancreatic, colon and lung cancers respectively [5,6]. Survival of the cancer cells that have mutated and activated oncogenes is dependent on the permanent presence and expression of these oncogenes; this phenomenon is called oncogene addiction. In the consequence of this phenomenon, inhibition of an active oncogene, leads to reverse cell transformation or induce cell death [7]. In spite of the fact that Ras oncogenes are important in the onset of cancer, there is no molecular targeted therapy drug to control the expression of this protein at present. This pharmacologic flaw may be because of the lack of suitable ligand binding pocket on the surface of Ras protein [8]. Also, the farnesyl transferase inhibitors, which inhibit the positioning of Ras in cytoplasmic side of membrane, have proved to be a failure in clinical trials [9]. Moreover, despite the absence of approved Ras-inhibiting therapeutics, some considerable efforts have been carried out in this regard. Some suitable inhibitors, having the capability of preventing interactions between Ras and upstream/downstream effectors; such as SOS and RAF kinases have previously been identified [10–13]. On the other hand, a number of successful good efforts have been exercised by using natural products to identify potential anticancer compounds [14]. Furthermore, wide innate diversity of natural products encourage the researchers to use high-throughput screening (HTS) methods to identify suitable inhibitors among them [15].

**Figure 1.**
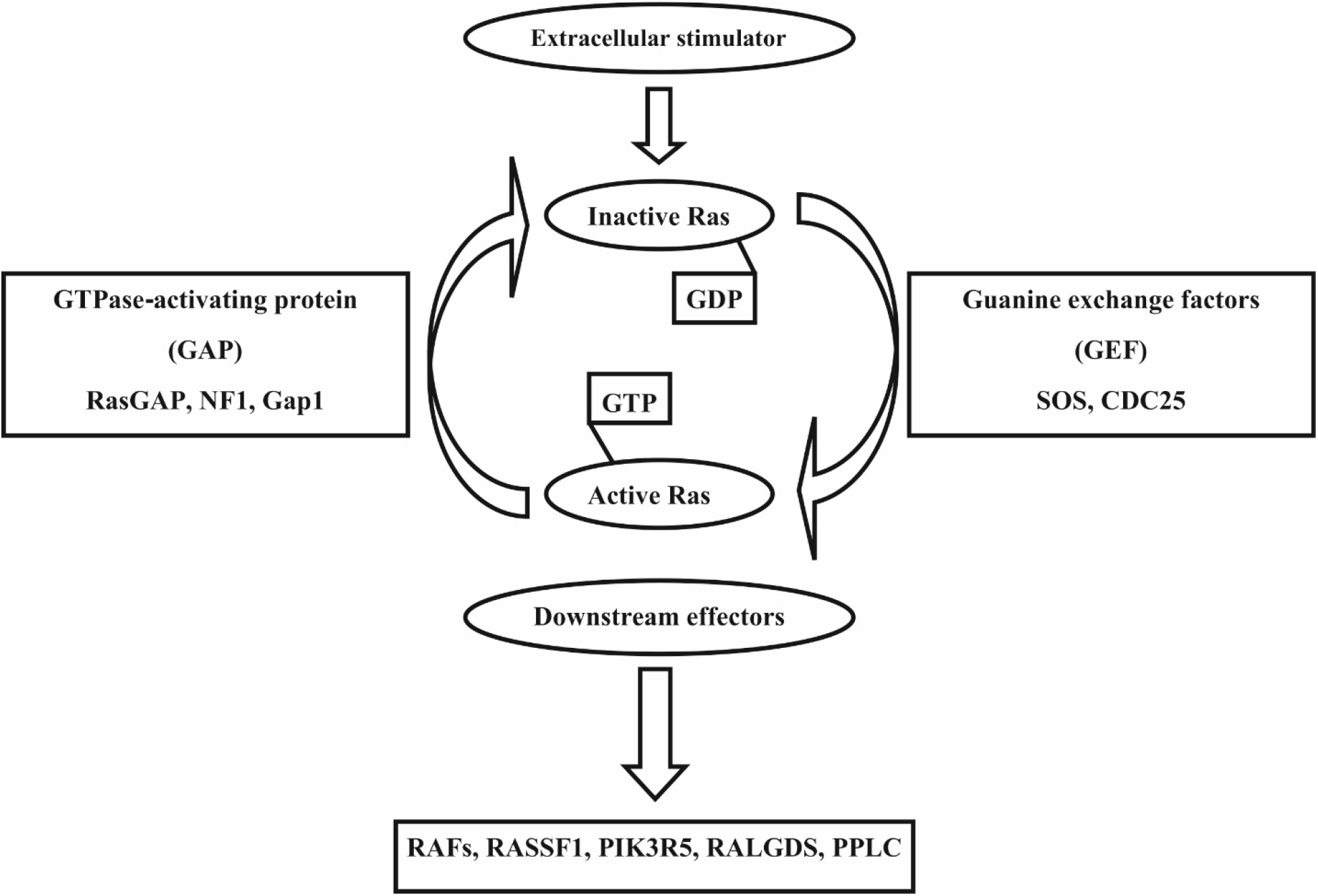
Master Switch function of Ras

Due to the fact that drug discovery is a highly expensive filed, using computational science such as computer-aided drug design or high throughput virtual screening (HTVS) has become attractive approach. Here, we employed the HTVS to identify new potent K-Ras^G12V^ inhibitors. HTVS has been done through Lipinski’s rule of 5 (RO5) for drug likeness screening, molecular docking method and binding free energy calculations, using MM-P/GBSA method and based on molecular dynamics (MD) simulation calculations. Hit ligands were identified via HTVS simulation. Then post MD simulation analysis of Hit ligands was performed to investigate the stability and interaction properties of Hit ligand-protein complexes.

## Materials and Methods

### Data collection

First, a 3D structure library was built for virtual screening, containing the molecular structures of 81481 small natural molecules from five subsets (MdpiNP, TcmNP, IBSNP, AcdiscNP and AmbintNP) of the ZINC database [16] in MOL2 format and the 3D structure of K-Ras (PDB ID 4EPY) [13] was obtained via RCSB database [17]. This structure had no missing loops or residues, had the resolution of 1.8 Å and contained a specific mutation (G12V), which was the main target of this study [13]. The ligand structures were taken from ZINC database in MOL2 format and were ready to Dock.

### Molecular dynamics studies

As a consequence of the specific situation of X-Ray crystallography, hydrogen atoms are missed and also detected protein conformation have minor differences with water-solved protein. Thus, for obtaining accurate results, we should fix these issues with proper settings. This method keeps reasonable balance between cost and accuracy. GROMACS 5 was used to carry out MD simulation, [18]. Ligand atoms were isolated from protein structure by UCSF chimera [19]. Then, the topology files of remaining structure were created by pdb2gmx application from GROMACS package using AMBER03 force field [20] and TIP3P water mode [21]. The topology parameters of ligands were calculated by ACPYPE [22], which uses Antechamber software [23].

The simulation was performed under periodic boundary condition (PBC) and the simulation box was cubic shape with minimal distance of 1.0 Å between protein and the edge of the box. The simulation box contained protein structure and hetero atoms of ligands, and was filled up by TIP3P water mode. In order to make the system neutral, physiological concentration of NaCl (0.15 mM) was applied to the system. Energy minimization of all atoms was done using steepest descent minimization algorithm and the maximum force was lessened to 1000 Kcal mol^−1^ nm^−1^. Then equilibration of each system was performed under 100ps NVT and 300ps NPT with position restraint for K-Ras^G12V^ and small molecules. As final step, 30ns MD simulation was performed with a time step of 2 fs at constant pressure (1 atm) and temperature (310 k). During MD simulation, PME [24], LINCS [25] and SETILE [26] algorithms were used to calculate long range electrostatic interactions, constrain the bond length and water molecule respectively. Moreover, velocity rescaling thermostat [27] and Parrinello-Rahman [28] barostat were used to couple temperature and pressure respectively. MD simulation information, including energy and trajectory information was saved every 2ps for further study and was finally analyzed by GROMACS utilities, UCSF chimera and LigPlot^+^ [29].

### Virtual high throughput screening (VHTS)

After primary preparations, our input data (ligands and protein structure files) was made available to initiate virtual screening. In the study ahead, virtual screening was performed in three steps including primary screening of drug-like compounds, molecular docking and binding free energy which was based on MD calculations. The study was carried out, based on the RO5 [30], using evaluator program from ChemAXON package[31]. Ligands library was screened and drug-like compounds were separated from other structures. Then, binding affinity of drug-like compounds to the binding site of K-Ras was computed using molecular docking. Molecular docking is an *in silico* method for finding the position of ligands onto its receptor. Our molecular docking calculation was performed by AutoDock Vina [32] that uses iterated local search global optimizer search algorithm and semi-empirical scoring function. Grid box and grid spacing was considered 25*25*25 and 1 Å, respectively. Grid box was centered on a known binding pocket and included alpha-helix-2 and beta-sheets-1, 2 and 3 that are significantly important in the formation of binding pocket and ligand binding [13]. Atomic re-ordering and Gasteiger charges were calculated by AutoDock Tools-1.5.6rc3 (ADT) [33] and Raccoon programs [34], respectively. Further, we assumed that during calculation, the ligand and protein structures are flexible and rigid, respectively. Then, the virtual screening was performed by VSDK program [35], and the results were sorted on the basis of ligand affinity and the ligands with high affinity were selected as appropriate docking results. Afterwards, the poses obtained from docking calculation were re-scored by X-Score [36] and Hit ligands were then determined. This procedure was validated with AutoDock Vina by comparing the docked poses and co-crystal ligand pose.

Finally, after Hit ligands determination by molecular docking and re-scoring, the ligand-protein complexes were simulated within 10 ns as described above and the results were subjected binding free energy calculations using MM-P/GBSA methods that demonstrates the role of electrostatic, van der Waals, polar solvation energies and solvent-accessible surface area (SASA) in ligand-protein binding and measured contribution of each residue in binding process, during MD simulation with the help of g-mmpbsa package [37]. A total of 100 snapshots were extracted from trajectory files, while the solvent molecules including water and ions were removed and binding free energy of each complex was calculated. Snapshot extraction was unbiased and based on equitable distribution of snapshots during simulation time. At the end, hit ligands were sorted base on their binding free energies and final selection was accordingly.

Furthermore, in order to gain broader understanding of MD Simulation calculations, post simulation analysis (RMSD and RMSF values and hydrogen bonds formation between ligand and protein over the simulation time) of Hit ligands were performed using GROMACS utilities. The last frame of trajectory file has been extracted and hydrogen bond and non-bound contacts, between ligands and protein structure were analyzed using UCSF chimera and LigPlot.

## Results and discussion

### Primary preparation

The compounds obtained from ZINC database, were ready-to-dock and required no additional preparations. But the protein was prepared by MD simulation and after 30 ns, the protein structure was equilibrated and hydrogen orientations were corrected. This result was confirmed by RMSD plot and the average RMSD during simulation was 0.152 nm (Fig5). Furthermore, RMSF data shows strong correlation of these findings with experimental results and demonstrated high RMSF value for Switch-2 region that confirms the role of Switch-2 fluctuations in binding pocket formation [13].

### Virtual Screening and re-scoring

As a result, 27684 ligands passed through RO5 principles and 52143 ligands were rejected. The ligands, which passed Lipinski RO5, were docked into the simulated structure of K-Ras simulated structure, re-scored by X-Score and 9 ligands (ZINC03845308, ZINC85592862, ZINC08876650, ZINC12889550, ZINC12889473, ZINC38139486, ZINC15671852, ZINC85567582 and ZINC03616630) were selected as Hit ligands. The computed affinity and average X-Score of selected compounds and reference ligands are shown in Table 1. In the present study, reference ligand was *(S) - N-(2-((1 H-indol-3-yl) methyl)-1 H-benzo[d] imidazol-5-yl) pyrrolidine-2-carboxamide* (Fig2A) that was co-crystalized with K-Ras structure (4EPY, PDB-ID) [13]. The affinity and average X-Score of the reference ligand to the protein binding pocket were calculated to be −8.5 Kcal/mol and 5.36, respectively. These results were considered as a threshold to judge molecular docking and re-scoring results. This ligand was observed to contribute in 2 H-bond and 34 hydrophobic interactions with protein binding pocket. Re-scoring by X-Score and comparing to with reference ligand let to the elimination of false-positive results.

**Figure 2.**
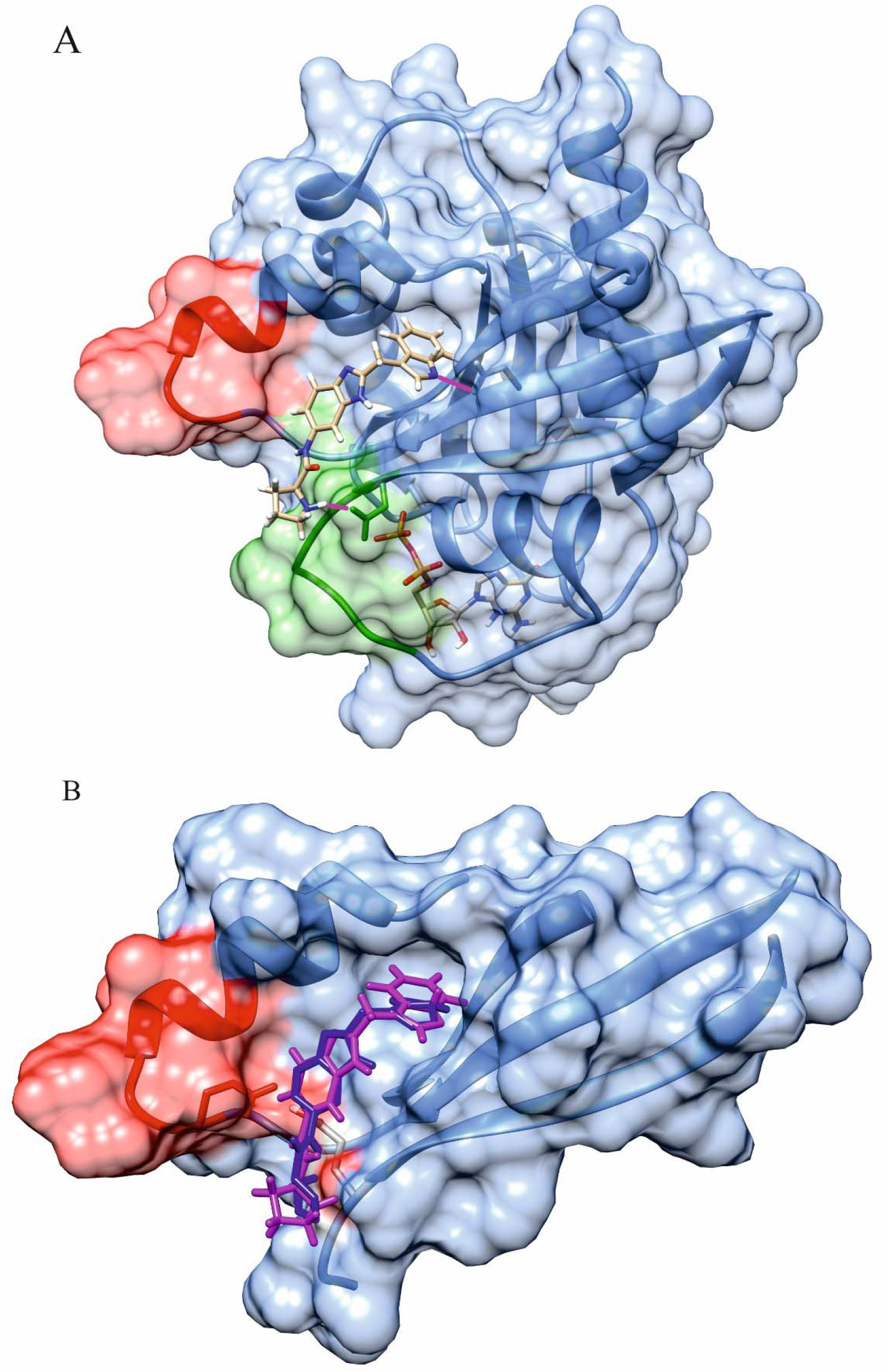
A) Whole structure of K-Ras protein. Switch-1 and switch-2 are colored forest green and red respectively. GDP and reference ligand are shown in atom stick form. B) Re-docked ligand was colored blue and crystallographic ligand is shown with magenta. Alpha helix-2 and beta sheets 1-3 were separated from protein. Also, the switch-2 is colored in red

**Table 1.** Calculation of docking and re-scoring score of 9 Hit ligands and reference ligand

| Ligand ID | affinity (kcal/mol) By AutoDock Vina | X-Score |  |  |  |
| --- | --- | --- | --- | --- | --- |
|  |  | HPScore | HMScore | HSScore | Average |
| **Reference Ligand** | -8.5 | 5.56 | 5.28 | 5.24 | 5.36 |
| **ZINC03616630** | -7.5 | 5.57 | 6.06 | 5.31 | 5.65 |
| **ZINC03845308** | -8.3 | 6.59 | 7.75 | 6.27 | 6.87 |
| **ZINC08876650** | -7.7 | 5.88 | 6.98 | 5.84 | 6.23 |
| **ZINC12889473** | -7.5 | 5.67 | 7.1 | 5.75 | 6.17 |
| **ZINC12889550** | -7.5 | 5.69 | 7.15 | 5.77 | 6.2 |
| **ZINC15671852** | -8.3 | 5.78 | 6.31 | 5.51 | 5.86 |
| **ZINC38139486** | -7.9 | 5.44 | 6.85 | 5.45 | 5.91 |
| **ZINC85567582** | -7.6 | 5.86 | 6.25 | 5.18 | 5.76 |
| **ZINC85592862** | -7.9 | 6.34 | 6.64 | 5.75 | 6.24 |

Validity ascertain of molecular docking procedure was confirmed via superimposition of the re-docked reference ligand with the ligand’s X-Ray crystallographic structure. Calculated RMSD value between crystallographic structure and re-docked ligand was 0.738 nm which is indicative of the accuracy of docking method (Fig2B).

### Binding free energy and post simulation analysis

Hit and reference ligand complexes were simulated for 10 ns as described above and the results were subjected to binding free energy calculations by MM-P/GBSA method. According to MM-PBSA results (Table 2), in the consequence of hydrophilic nature of reference and Hit ligands and shallowness of binding pocket, polar solvation energy of all ligands was observed to have positive values. Also, SASA energy of Hit and reference ligands were non-significant. On the other hand, data analysis of calculated values of van der Waals and electrostatic energy reflect the type of balance between the role of van der Waals and electrostatic interactions in ligand binding. In this case, if the sum of van der Waals and electrostatic energy is a negative value and overcome the positive value of the polar solvation energy, binding free energy could become negative and we can assume the formation of ligand-protein complex, which is a thermodynamically possible reaction. Binding free energies of 4 ligands (ZINC15671852, ZINC85592862, ZINC85567582 and ZINC03616630) were less than that of the reference ligand, were selected as final Hit ligands and were considered for further studies. Reference ligand and ZINC15671852 have partially similar binding pattern and were more affected by van der Waals energy than electrostatic energy. Although, binding of ZINC85592862, ZINC85567582 and ZINC03616630 to the protein are under the influence of electrostatic energy instead of van der Waals energy. Also, using the MM-GBSA method, binding free energies of ZINC15671852, ZINC85592862, ZINC85567582, ZINC03616630 and reference ligand were decomposed and energy contribution of each residue was predicted (XVG files of MM-GBSA calculations are provided in attachment). MM-GBSA results characterized the binding patterns of ligands (final Hit and reference ligand) and binding pocket residues (1-80 which include beta sheet 1-3 and alpha helix 2) and specified critical residues in binding pocket. Energy contribution of binding pocket residues in ligand-protein interaction showed in Fig3. Also, energy contribution value of critical residues of each complex was determined (Table 3). According to the MM-GBSA results GLU4, ASP34, GLU38, ASP39, ASP58, GLU63, GLU64 and ASP70 residues may have critical role and effective contribution in binding of final Hits. Furthermore, energy contribution of binding pocket residues, reveals that ZINC85592862, ZINC85567582 and ZINC03616630 may have similar binding pattern with the reference ligand.

**Figure 3.**
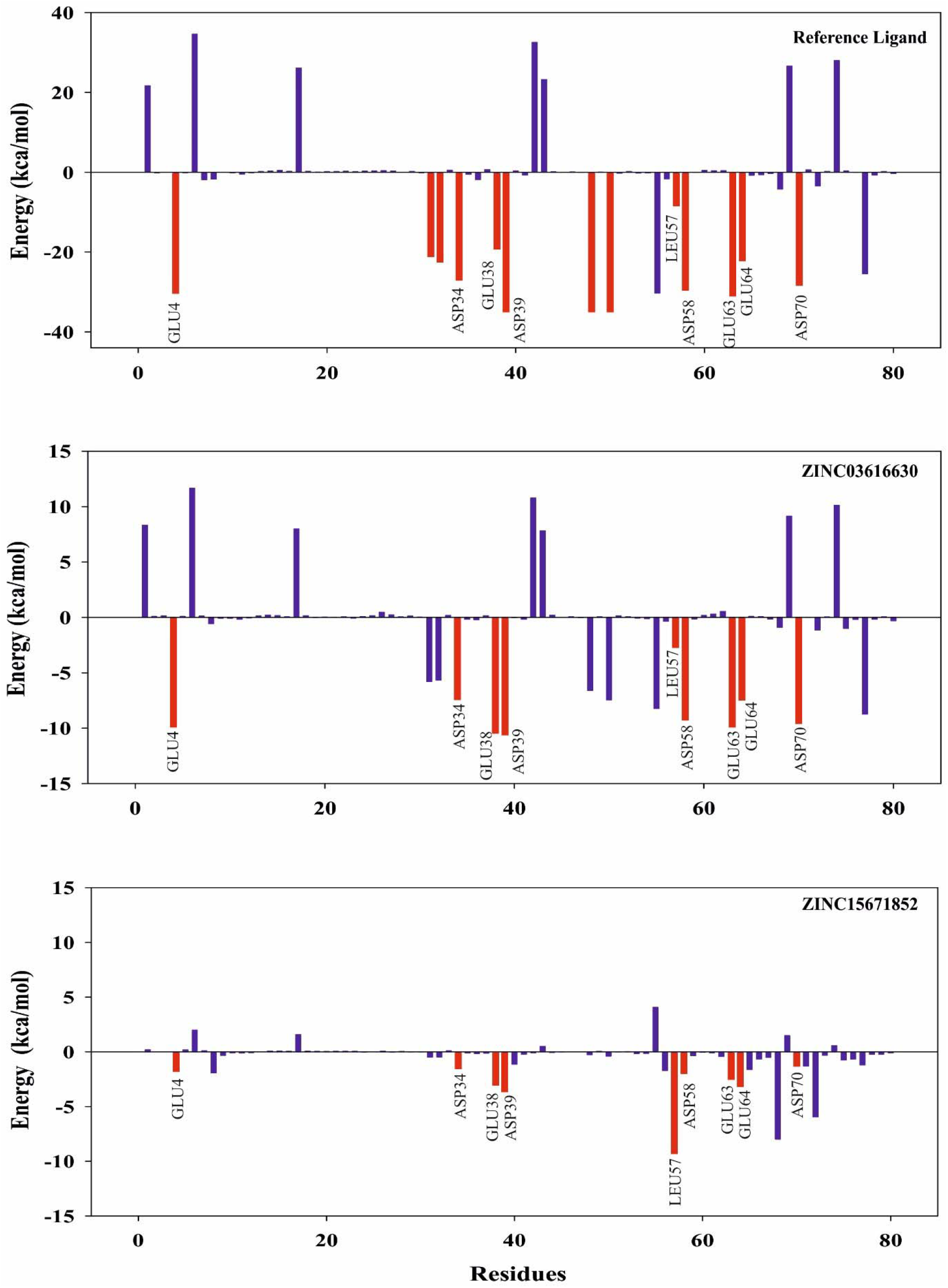
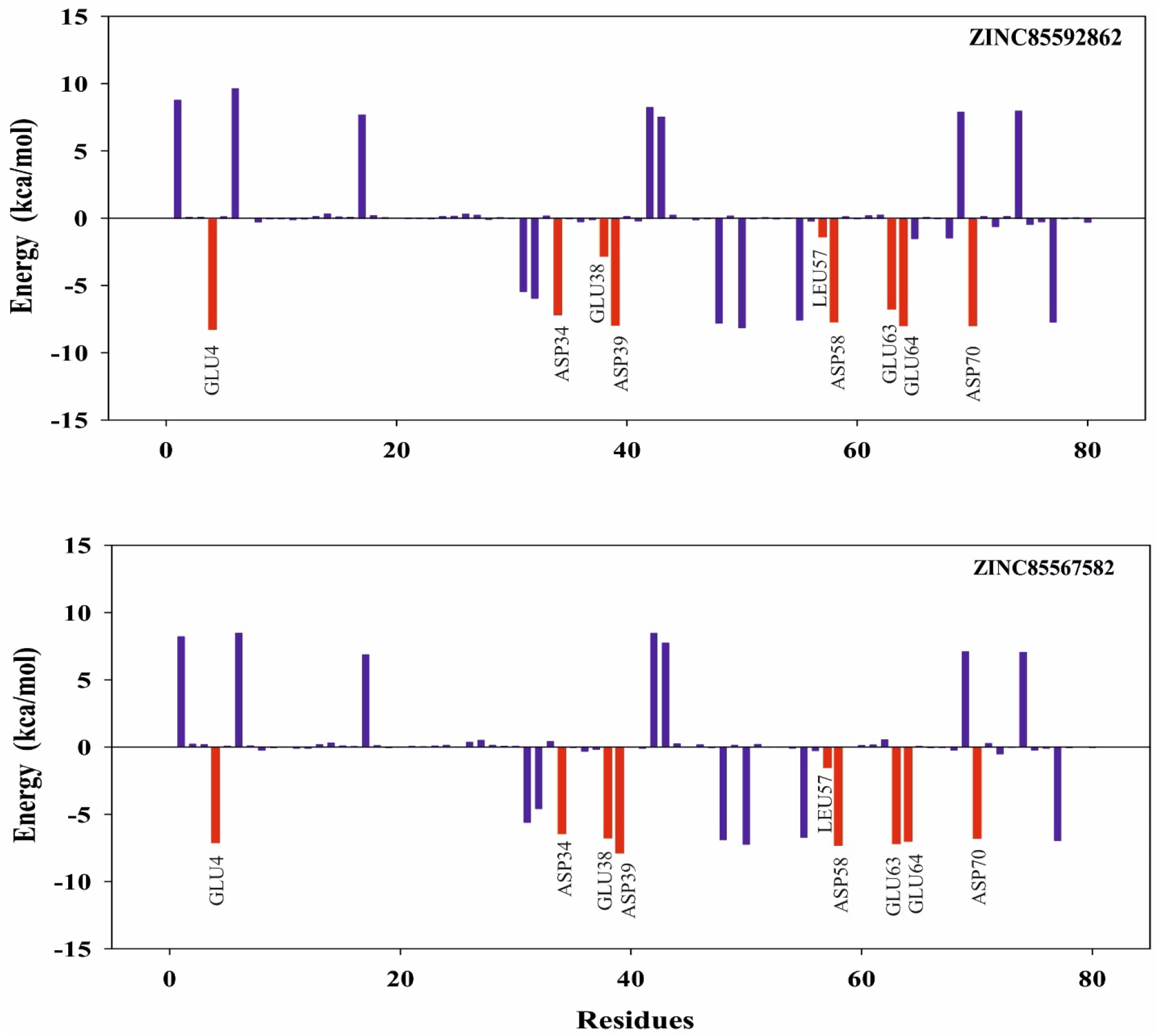
Energy contribution of binding pocket residues in ligand-protein interaction. Consensus residues between final Hit and reference ligand are shown in red color

**Table 2.**
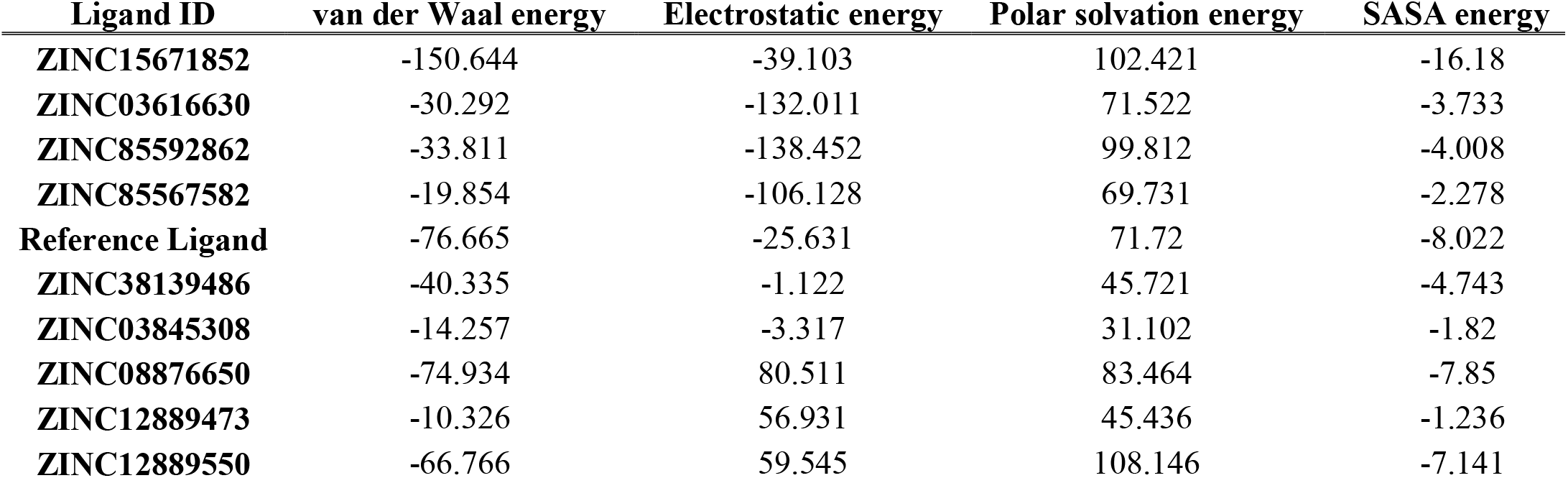
MM-PBSA calculated energies for 9 Hit ligands and reference ligand

**Table 3.**
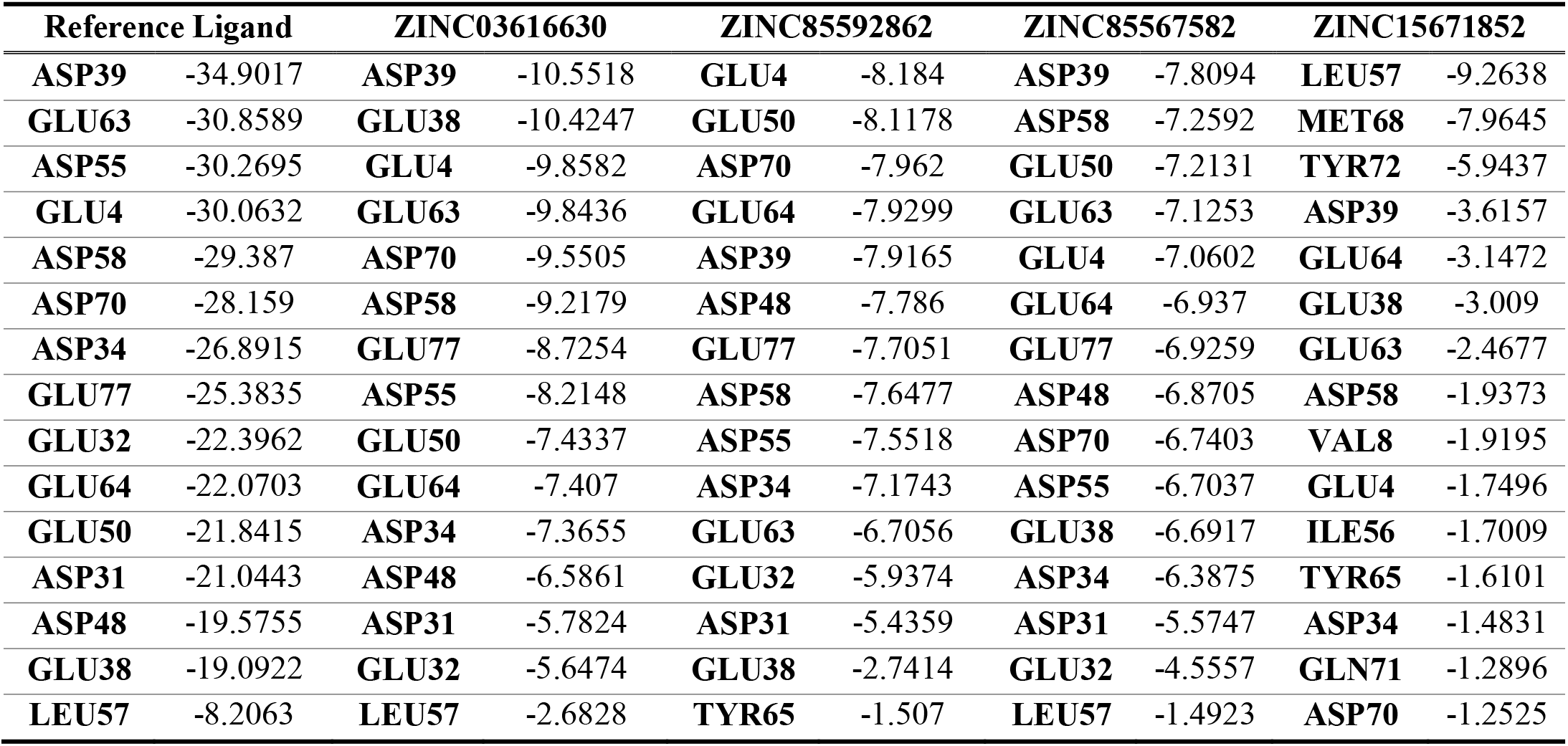
MM-GBSA calculated energy contribution of critical residues of reference and final Hit ligands

As the Ras protein was simulated with the new ligands and the conformation changes in residue side chain at binding pocket was carried out in the present study, the last frame of Hits and reference ligand complexes were extracted from trajectory. Hydrogen bonds and non-bonded contacts were diagnosed by LigPlot+ and chimera respectively (LigPlot+ data are shown in supplementary, Tables 1S and 2S). Also, Hit and reference ligand’s interactions with protein binding site were shown (Fig4 and 1S). RMSD value is the primary standard to evaluate the stability of protein structure. Also, the results shown ligand-protein complexes were relatively stabilized after 5 ns MD simulation (Fig5A). Average RMSD values of Hit and reference ligands ranged from 0.126 to 0.152 nm (XVG files of RMSD calculations are provided in attachment).

**Figure 4.**
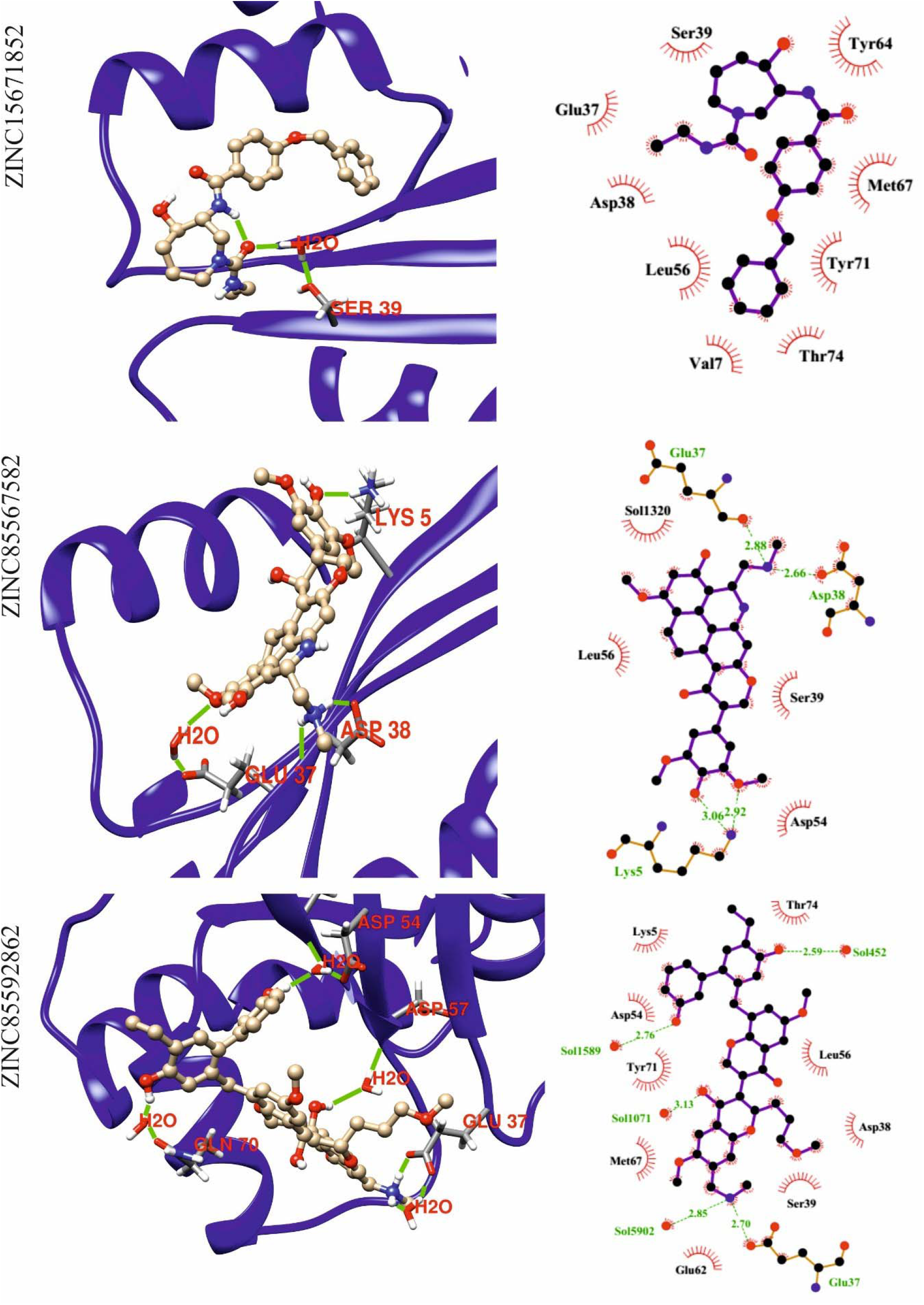
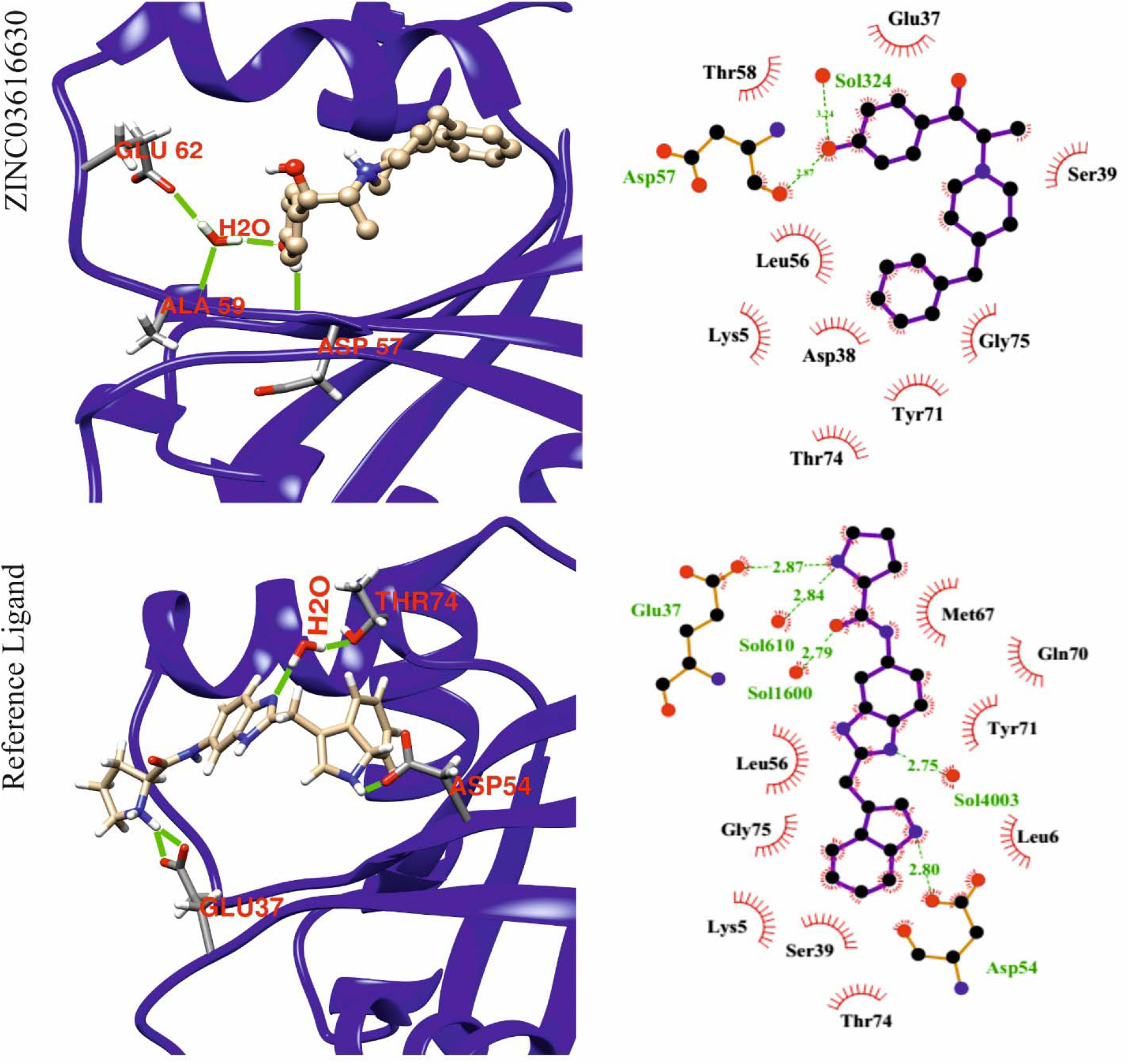
Post simulation analysis of Hit ligands and protein binding pocket with chimera

**Figure 5.**
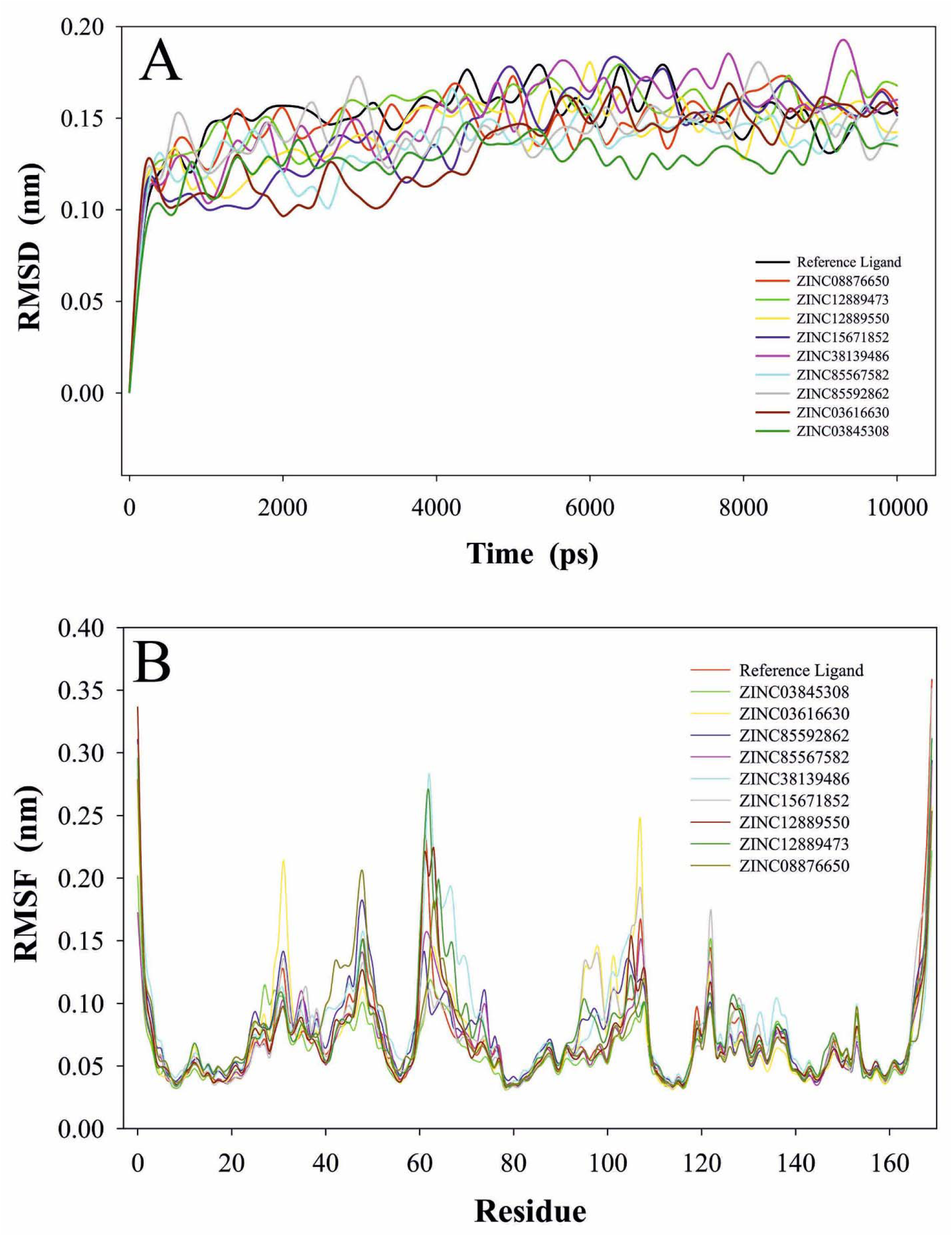
A) Root Mean Square Deviation of protein C-alpha during 10 ns simulation with the ligands, B) Root Mean Square Fluctuations of CA as a function of residue number for all the simulations. Different colors represent different trajectories of Ras protein in the present and absent of ligands

Furthermore, to evaluate the effect of ligand binding on the flexibility of each residue during MD simulation, RMSF values of ligand-protein complexes and apo-protein were calculated accordingly (Fig5B). Analysis of the present results have revealed that the binding of Hit and reference ligands to this pocket, had no effect on Switch-1 fluctuation, but Switch-2 fluctuation was affected by ligand binding (XVG files of RMSF calculations are provided in attachment). Furthermore, H-bond formation between Hit or reference ligands with the K-Ras protein are calculated and shown in Fig6 (XVG files are provided in attachment).

**Figure 6.**
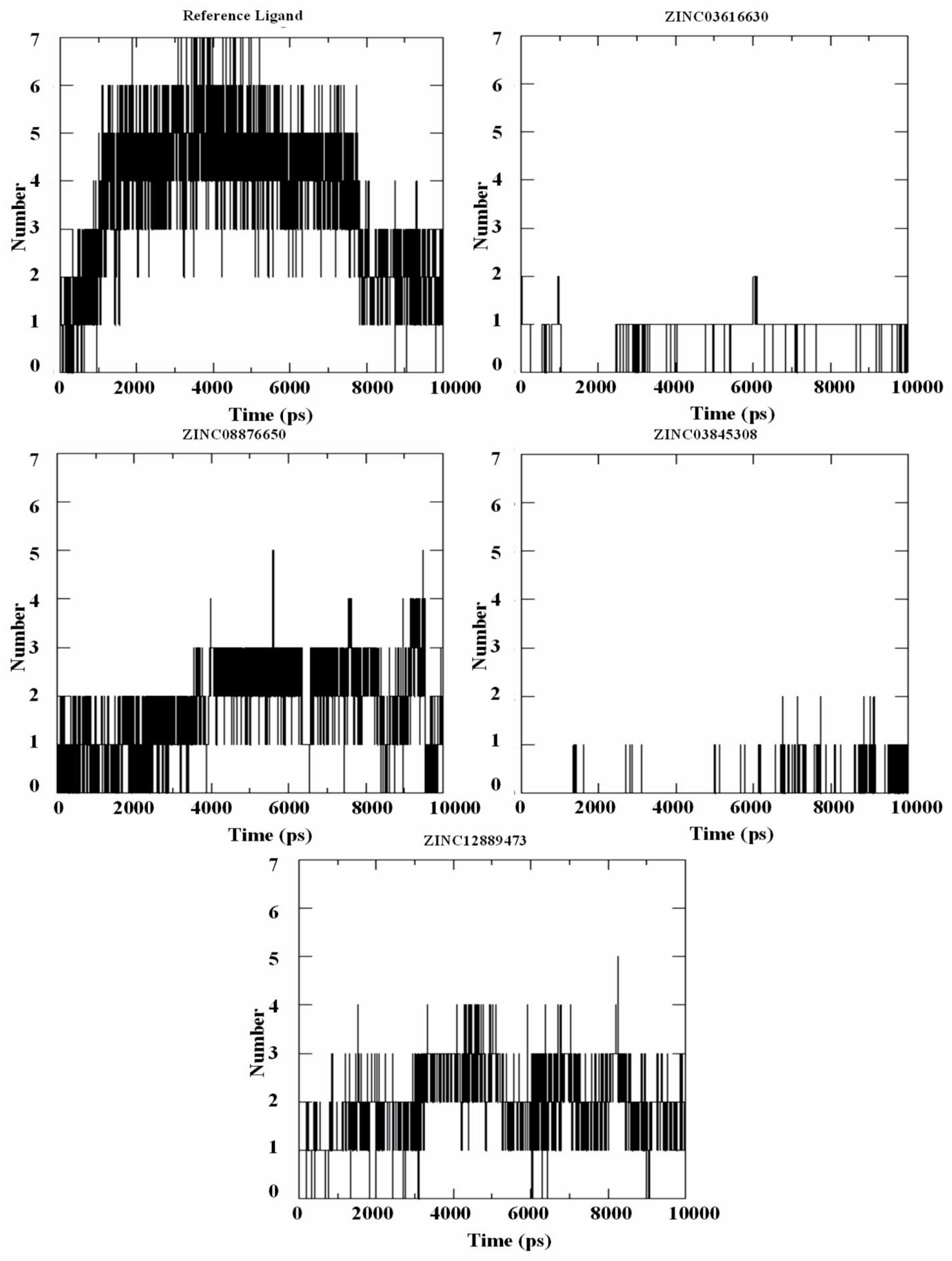
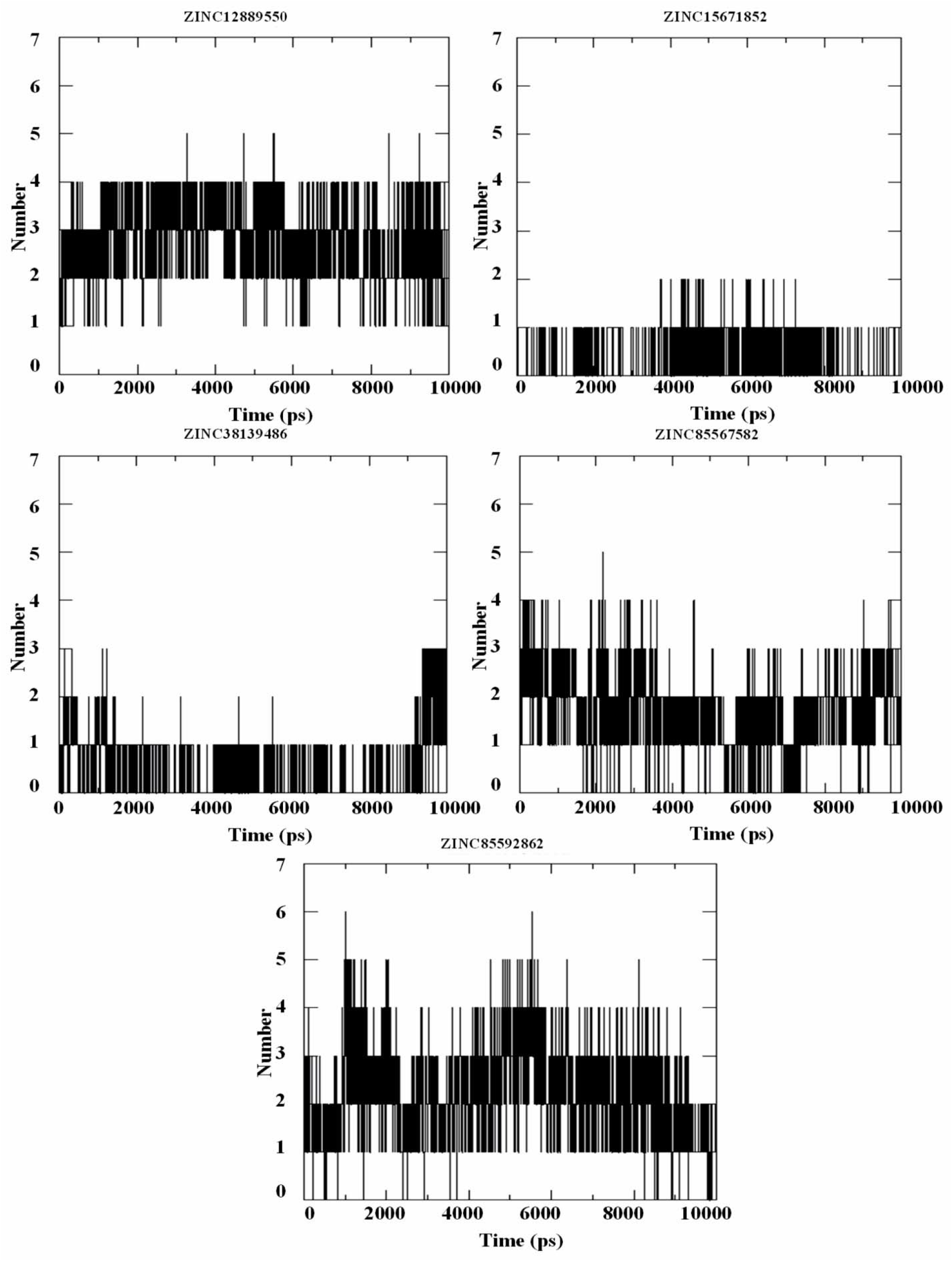
Hydrogen bond formation in ligand-protein complex during 10 ns MD simulation

## Conclusions

During the present study, VS was carried out using AutoDock Vina and X-Score re-soring program to identify new binders for K-Ras^G12V^ among three natural product categories from ZINC database. Subsequently ZINC03845308, ZINC85592862, ZINC08876650, ZINC12889550, ZINC12889473, ZINC38139486, ZINC15671852, ZINC85567582 and ZINC03616630 were selected as Hit ligands and subjected to MD simulation calculations. The RMSD analysis demonstrated the complex of these small molecule with the K-Ras protein are relatively stable. then free binding energy calculations is applied and finally, four Hits (ZINC15671852, ZINC85592862, ZINC85567582 and ZINC03616630) were identified as potential inhibitors for K-Ras^G12V^. Negative binding free energy of these Hits reveals relatively stable affinity of them to the binding pocket of K-Ras^G12V^ protein. Also, energy contribution analysis of binding pocket residues shows that possible binding behavior of ZINC85592862, ZINC85567582 and ZINC03616630 are similar to the reference ligand.

## Acknowledgments

This investigation was supported by a grant from Department of Biology, Faculty of Science, Golestan University, Gorgan, Iran.

## Supplementary

**Figure 1S.**
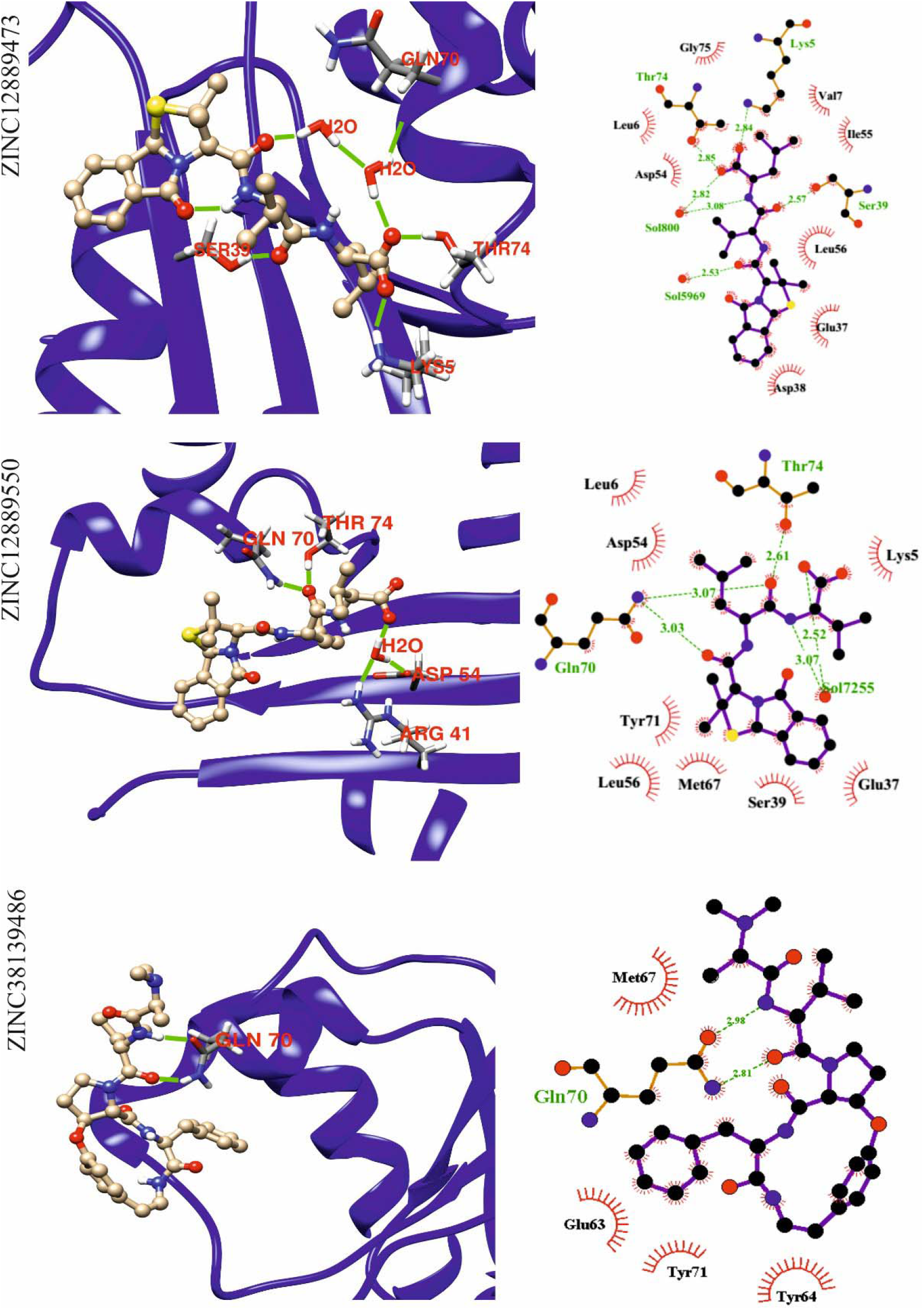
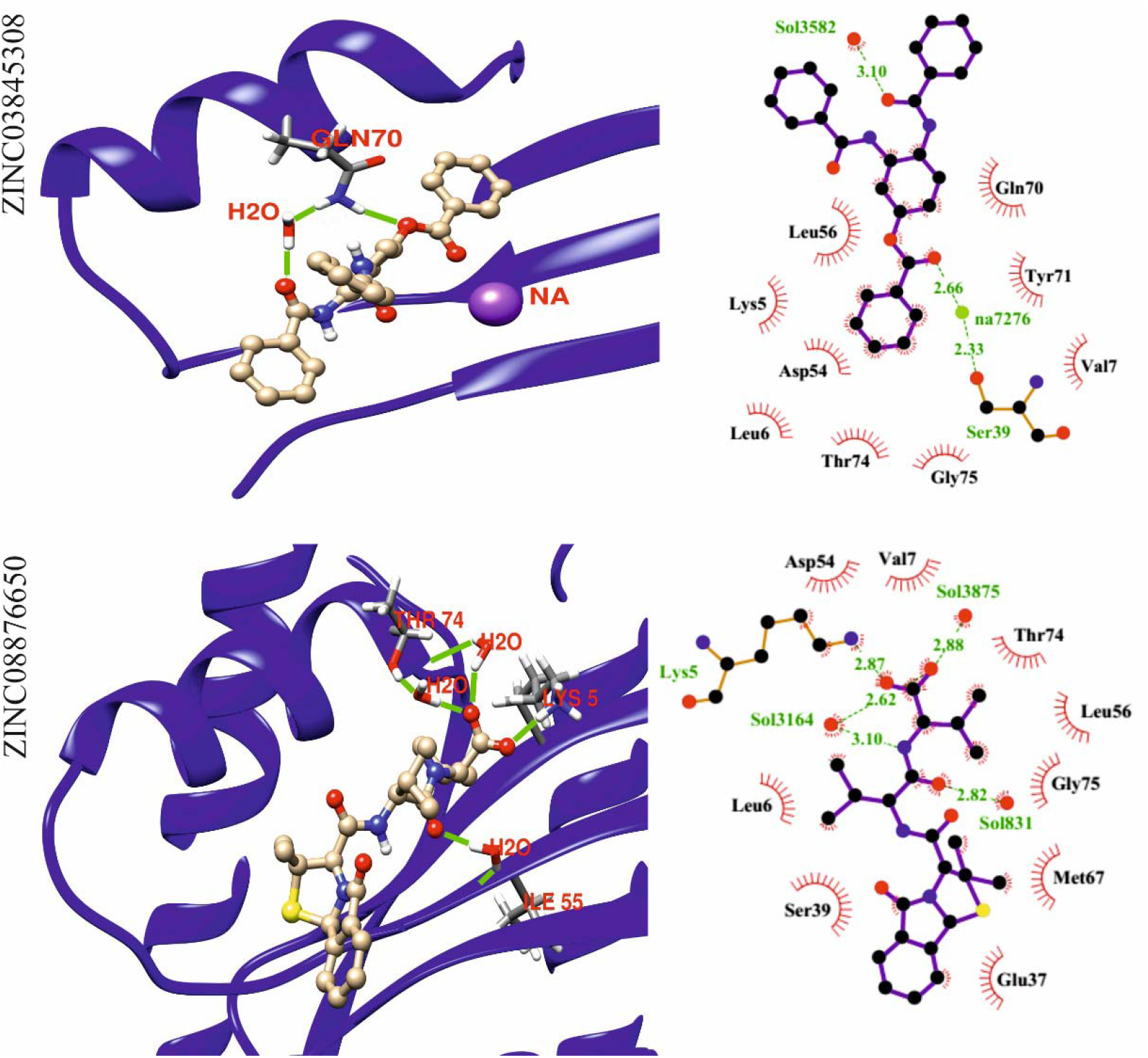
Post simulation analysis of Hit ligands and protein binding pocket with chimera

**Table 2S.** Post simulation hydrogen bonds of Hit compounds with the protein binding pocket

| Ligand code | Donor |  | Acceptor |  | Distance |
| --- | --- | --- | --- | --- | --- |
| **Reference Ligand** | LIG | N21 | OD2 | ASP-54 | 2.8 |
|  | LIG | N13 | OE2 | GLU-37 | 2.87 |
| **ZINC03616630** | LIG | O12 | O | ASP-57 | 2.87 |
| **ZINC03845308** | LIG | N9 | OW | SOL | 3.23 |
|  | NA | NA | OG | SER-39 | 2.33 |
|  | NA | NA | O27 | LIG | 2.66 |
|  | SOL | OW | NA | NA | 2.39 |
|  | SOL | OW | NA | NA | 2.47 |
|  | SOL | OW | NA | NA | 2.45 |
|  | SOL | OW | NA | NA | 2.42 |
| **ZINC08876650** | LYS-5 | NZ | O17 | LIG | 2.87 |
| **ZINC12889473** | NA | NA | OD1 | ASP-54 | 2.36 |
|  | NA | NA | O | ILE-55 | 2.27 |
|  | LIG | N17 | O28 | LIG | 2.73 |
|  | LYS-5 | NZ | O63 | LIG | 2.84 |
|  | NA | NA | O11 | LIG | 3.24 |
|  | SER-39 | OG | O11 | LIG | 2.57 |
|  | THR-74 | OG1 | O8 | LIG | 2.85 |
|  | NA | NA | OG | SER-39 | 2.49 |
|  | NA | NA | O | TYR-40 | 2.45 |
| **ZINC12889550** | GLN-70 | NE2 | O19 | LIG | 3.03 |
|  | GLN-70 | NE2 | O8 | LIG | 3.07 |
|  | NA | NA | OE2 | GLU-3 | 2.37 |
|  | NA | NA | O63 | LIG | 2.41 |
|  | NA | NA | O16 | LIG | 2.36 |
|  | SOL | OW | NA | NA | 3.33 |
|  | SOL | OW | NA | NA | 2.48 |
|  | THR-74 | OG1 | O8 | LIG | 2.61 |
| **ZINC38139486** | GLN-70 | NE2 | O13 | LIG | 2.81 |
|  | LIG | N6 | OE1 | GLN-70 | 2.98 |
| **ZINC85592862** | LIG | N41 | OE2 | GLU-37 | 2.7 |
| **ZINC85567582** | LIG | N2 | OD1 | ASP-38 | 2.66 |
|  | LIG | N2 | O | GLU-37 | 2.88 |
|  | LYS-5 | NZ | O35 | LIG | 2.92 |
|  | LYS-5 | NZ | O34 | LIG | 3.06 |

**Table 2S.** Post simulation non-bonded contacts of Hit compounds with the protein binding pocket

| Residue | Reference<br>Ligand | ZINC03616630 | ZINC03845308 | ZINC08876650 | ZINC12889473 | ZINC12889550 | ZINC15671852 | ZINC38139486 | ZINC85592862 | ZINC85567582 |
| --- | --- | --- | --- | --- | --- | --- | --- | --- | --- | --- |
| LYS5 | 4 | 2 | 2 | 5 | 2 | 3 |  |  | 6 | 6 |
| LEU6 | 1 |  | 1 | 3 | 3 | 1 |  |  |  |  |
| VAL7 |  |  | 2 | 2 | 1 |  | 1 |  |  |  |
| GLU37 | 3 | 5 |  | 1 | 3 | 1 | 3 |  | 8 | 3 |
| ASP38 |  | 1 |  |  | 8 |  | 1 |  | 2 | 9 |
| SER39 | 3 | 2 |  | 3 | 5 | 2 | 1 |  | 2 | 3 |
| ASP54 | 6 |  | 1 | 2 | 5 | 4 |  |  | 1 | 5 |
| ILE55 |  |  |  |  | 2 |  |  |  |  |  |
| LEU56 | 4 | 5 | 5 | 1 | 6 | 4 | 2 |  | 2 | 2 |
| ASP57 |  | 3 |  |  |  |  |  |  |  |  |
| THR58 |  | 2 |  |  |  |  |  |  |  |  |
| GLU62 |  |  |  |  |  |  |  |  | 3 |  |
| GLU63 |  |  |  |  |  |  |  | 8 |  |  |
| TYR64 |  |  |  |  |  |  | 2 | 12 |  |  |
| MET67 | 4 |  |  | 2 |  | 1 | 12 | 7 | 8 |  |
| GLN70 | 1 |  | 7 |  |  | 6 |  | 9 |  |  |
| TYR71 | 2 | 4 | 3 |  |  | 2 | 3 | 4 | 7 |  |
| THR74 | 5 | 1 | 5 | 1 | 6 | 3 | 1 |  | 7 |  |
| GLY75 | 1 | 2 | 1 | 1 | 2 |  |  |  |  |  |
| SUM | 34 | 27 | 27 | 21 | 43 | 27 | 26 | 40 | 46 | 28 |

